# Words Matter with Age: Instructions Dictate Self-Selected Walking Speed in Young and Older Adults

**DOI:** 10.1101/2025.07.18.665559

**Authors:** Kenneth Harrison, Sarah A. Brinkerhoff, Emma Frese, Brandon M. Peoples, Keven G. Santamaria Guzman, David T. Redden, Jaimie A. Roper

## Abstract

**Background:** Our previous work demonstrated that in young adults, 61% of gait speed variance was attributable to instruction type. However, no study has investigated whether verbal instructions differentially influence older adults. Research question: This study investigated how walking prompts contribute to gait speed variability across age groups. Methods: Thirty-four young adults (21±2 years) and twenty-eight older adults (70±5 years) performed walking trials responding to 24 different instructions. Results: Average walking speed was 1.23±0.30 m/s. Between-subject variance accounted for 25.3% of the total variance, while between subject and instruction variance components accounted for 76.1% of the total variance. When analyzed separately, variance due to instructions accounted for similar amounts of total variance within older adults (56.9%) and young adults (55.7%), a statistically non-significant difference (p = 0.85). A significant age-instruction interaction (χ^2^=76.84, df = 23. p<0.001) revealed that the age differences between average gait speed depended on which instruction was given. Complex instructions elicited the largest between group differences (β:−0.24 to −0.32 m/s), while simple tasks showed minimal differences (β:−0.03 to −0.06 m/s). Significance: Instructions explain similar variance within each age group (~56%), but the model treating instructions as fixed effects captures how different age groups respond to the same instructions. These findings highlight the critical importance of instruction standardization in gait assessment protocols, as systematic age-related differences in instruction interpretation can significantly impact measured outcomes. Instructions that produce minimal between-group differences may be most appropriate for standardized clinical assessments, while those showing larger age effects may be valuable for detecting age-related changes in gait control.

## Introduction

Healthy young adults (HYA) are often used as a standard against which to compare various populations, however the accuracy of these comparisons depends on both the consistency of data collection methods and the understanding of how assessment protocols might differentially affect various populations. Our previous work demonstrated that in young adults, 61% of gait speed variance was attributable to the specific instructions given, with average speeds ranging from 0.91 m/s to 2.16 m/s across 24 different prompts [1]. Gait speed is particularly important in older adult populations, where it serves as a key predictor of functional decline, falls risk, and mortality [2]. Understanding how verbal instructions influence gait speed in older adults is crucial for both research reliability and clinical assessment. Currently, the literature reports varying gait speeds for older adults, but these differences might be attributable to inconsistent instructions rather than true population differences.

Individuals may vary walking speeds based on everyday activities such as grocery shopping or walking to an appointment. However, the speeds of these everyday activities may vary not only between individuals but also between age groups. Previous research has shown that while walking speed generally declines with age [3], the extent to which instruction type influences this decline remains unclear. Understanding these potential age-related differences in response to verbal prompts could significantly impact how we assess and compare gait across the lifespan. The primary aim of this study was to investigate how verbal instructions influence gait speed variance in both healthy young and older adult volunteers. We provided individuals from both age groups with the same twenty-four walking instructions following the same predetermined order for all participants. The 24 walking instructions were selected to represent the range of verbal prompts commonly used across different gait assessment protocols in clinical and research settings. Currently, there is no universally standardized instruction for measuring self-selected walking speed, with clinicians and researchers employing various verbal cues ranging from simple directions (e.g., “walk at a comfortable speed”) to more specific prompts (e.g., “walk at your typical speed” or “walk as fast as possible”). These instruction variations may contribute to the inconsistent gait speed values reported across studies and clinical assessments, making it difficult to establish reliable normative values or compare results between different protocols. We hypothesized variability due to prompts in both age groups, with potentially greater variability in older adults. The second primary aim was to measure age-related changes in gait speed per instruction and provide reference values for clinicians. This information will be valuable for researchers and clinicians working with a wide range of age groups, allowing for more accurate comparisons and assessments of gait performance across the lifespan. Furthermore, understanding how different age groups interpret and respond to various walking instructions could lead to more standardized and effective gait assessment protocols.

## Methods

### Participants

34 young adults (age 21 ± 2 years) and 28 older adults (age 70 ± 5 years) recruited from the Auburn University community participated in this study. All participants were free of lower extremity bone fractures, muscle tears, and joint dislocations for at least six months and did not have any known issue that affected their typical walking.

### Procedures

All participants provided written informed consent before participating as approved by the University Institutional Review Board. APDM were attached using a 6-sensor set up (Lumbar, sternum, wrist, and ankle) and walking trials were performed within a 12.1-meter lab space free from distraction. The same research assistant provided 24 different instructions in the same predetermined order for all participants (Table 1). Participants were instructed to cross the lab 3 times per instruction. Gait cycle parameters were averaged across each instruction per participant. Walking speed was derived from the APDM Mobility Lab system using the validated algorithms for stride velocity calculation [4], [5]. The 6-sensor configuration (lumbar L5, sternum, bilateral wrists, bilateral ankles) has demonstrated strong concurrent validity with instrumented walkway systems (r > 0.95). Test-retest reliability for walking speed measurement has been reported as ICC > 0.90 in both young and older adults [6], [7]. For each trial, participants walked continuously across the 12.1-meter space three times without stopping. APDM Gait detection algorithms register gait cycles automatically, with delineated temporal buffers for gait initiation cycles.

**Table 1.** Instructions given to participants during gait testing with group mean and standard deviation across groups.

| Prompt | Young Mean and (SD)<br>m/s | Old Mean and (SD)<br>m/s |
| --- | --- | --- |
| 11. Walk as if you were walking in the grocery store. | 1.04 (0.20) | 0.83 (0.24) |
| 4. Walk across the walkway at a slow speed. | 0.82 (0.22) | 0.83 (0.16) |
| 23. Walk as if you were walking a small dog. | 1.18 (0.17) | 0.86 (0.19) |
| 5. Walk as if you were walking through a park. | 1.09 (0.19) | 0.89 (0.20) |
| 14. Walk as if you were walking home after a really bad day. | 1.06 (0.20) | 0.93 (0.22) |
| 12. Walk as if you were walking home with a group of friends at night. | 1.13 (0.17) | 0.92 (0.20) |
| 21. Walk as if you were walking through a field. | 1.19 (0.16) | 1.00 (0.19) |
| 9. Walk as if you were walking a big dog. | 1.27 (0.20) | 1.06 (0.22) |
| 3. Walk as if you were walking from the bed to the bathroom at night. | 1.11 (0.22) | 1.07 (0.18) |
| 7. Walk away from the table. | 1.27 (0.20) | 1.08 (0.12) |
| 1. Walk at a comfortable speed. | 1.15 (0.14) | 1.12 (0.15) |
| 17. Walk at your own self-selected speed. | 1.27 (0.21) | 1.16 (0.18) |
| 6. Walk as if you were walking home alone at night. | 1.33 (0.23) | 1.17 (0.20) |
| 13. Walk towards the table. | 1.33 (0.17) | 1.17 (0.13) |
| 24. Walk as if you were going to pick an item up off of a table. | 1.34 (0.18) | 1.18 (0.15) |
| 8. Walk at the speed in which you would walk on a busy sidewalk. | 1.36 (0.19) | 1.21 (0.19) |
| 20. Walk across the walkway at your typical speed. | 1.33 (0.22) | 1.22 (0.19) |
| 10. Walk across the walkway as if you're trying to keep up with kids. | 1.48 (0.19) | 1.32 (0.17) |
| 22. Walk as if you were walking across the street. | 1.36 (0.18) | 1.34 (0.19) |
| 15. Walk as if you were walking to catch a bus or taxi. | 1.59 (0.20) | 1.41 (0.22) |
| 18. Walk as if you were walking jay-walking. | 1.50 (0.19) | 1.38 (0.18) |
| 2. Walk across the walkway in under 20 seconds. | 1.48 (0.22) | 1.44 (0.21) |
| 16. Walk as if you were walking to a meeting and you were late. | 1.63 (0.16) | 1.48 (0.20) |
| 19. Walk as fast as possible. | 1.88 (0.23) | 1.62 (0.24) |

### Data analysis

All statistical analyses and visualizations were conducted in R studio (Posit Software, version 2023.12.0, PBC, Build 369) using R (version 4.3.1). Our statistical approach addressed two primary aims. We first estimated an unconditional model with only a random intercept for subject to determine the proportion of variance explained by individual differences. Next, we added a random intercept for instruction type to quantify the additional variance explained by different walking prompts across both age groups combined. Models were compared using likelihood ratio tests, Akaike information criterion (AIC), and Bayesian information criterion (BIC), with lower values suggesting better fit. Homogeneity of variance between age groups was assessed using Levene’s tests for each instruction. Statistical significance was set at p < 0.05.

For our primary aim we examined how much variance in gait speed was explained by instructions within each age group separately. For both young and older adults, we calculated intraclass correlation coefficients (ICC) to determine the proportion of variance attributable to different walking prompts versus individual differences. We compared these group-specific ICCs using bootstrapping methods (5000 simulations) to determine if the influence of instructions on walking speed varied between age groups. For our second aim, we employed an interaction model to examine age-related differences in response to specific instructions. We fitted a model including an interaction between age group and instruction type, which allowed us to identify which prompts elicited similar or different responses across age groups. This interaction model produced coefficients (β values) that quantified the magnitude of age-related differences for each specific instruction, while still accounting for individual variation through random subject effects.

## Results

### Participant Demographics

A total of 62 individuals participated in this study: 34 young adults (age 21±2 years, range 18-30 years) and 28 older adults (age 70±5 years, range 64-81 years). The young adult group consisted of 24 females and 10 males, while the older adult group included 14 females and 14 males. All participants were community-dwelling, independently mobile without assistive devices, and reported being generally healthy. Young adults were primarily university students, while older adults were local community members. All participants completed the full testing protocol with no adverse events.

### Gait speed

Over all 24 prompts, average walking speed across both age groups was 1.23 ± 0.30 m/s (Figure 1). As gait speed cannot be zero or less, but can be any reasonable positive number, the data are bounded on the left and not bounded on the right. The data are non-normal and positively skewed, which can be expected given the bounds of the data. Table 1 reports the means and standard deviations for each walking prompt across groups, revealing substantial variation in gait speed depending on the specific instruction provided, with speeds ranging from 0.83 m/s for both groups at slow speeds to maximum values of 1.88 m/s for young adults and 1.62 m/s for older adults when walking as fast as possible (Figure 2).

**Fig. 1.**
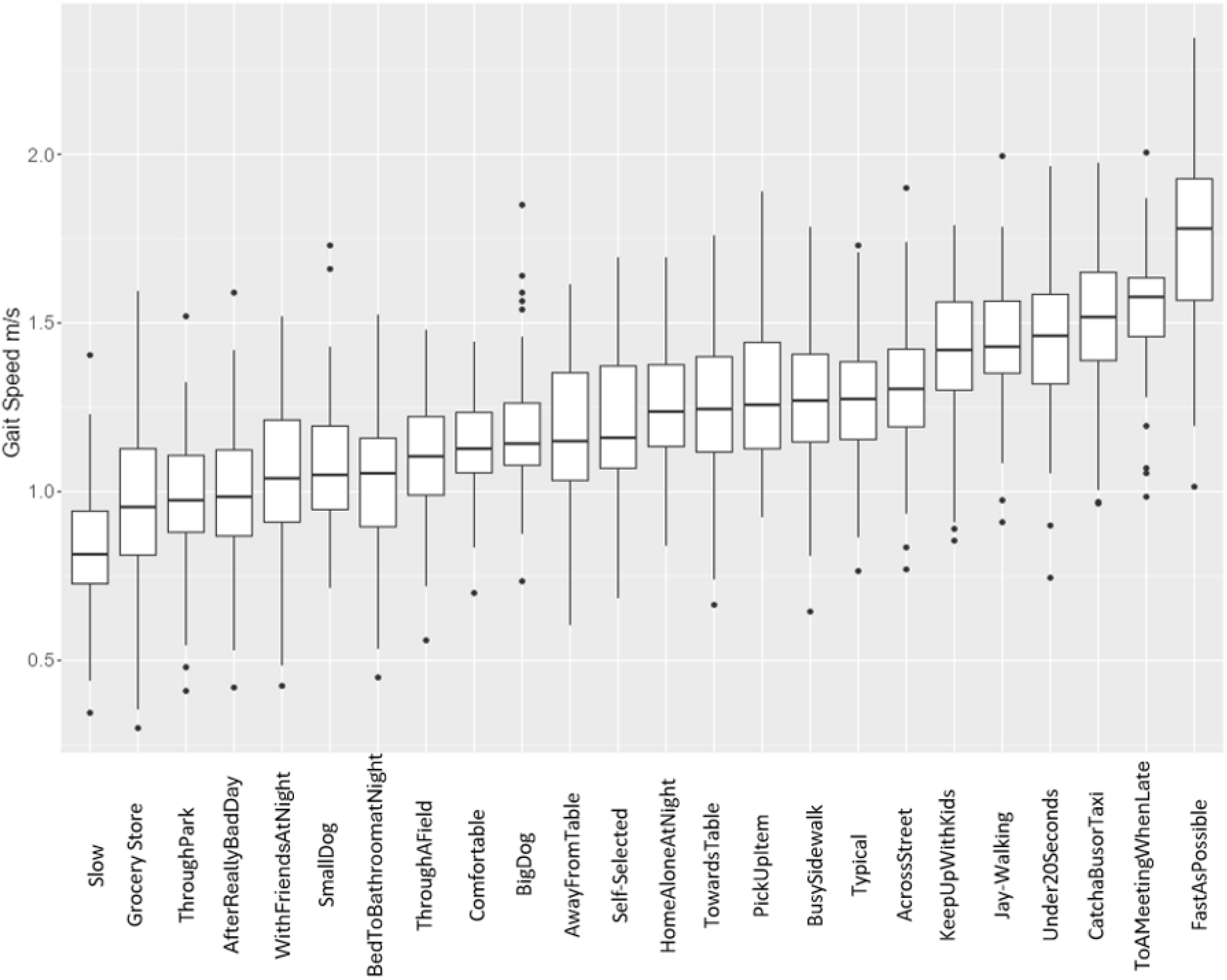
Distribution of speed by prompt ordered from slowest to fastest median speed. The box plot displays minimum, first quartile, median, third quartile, and maximum speeds.

**Fig. 2.**
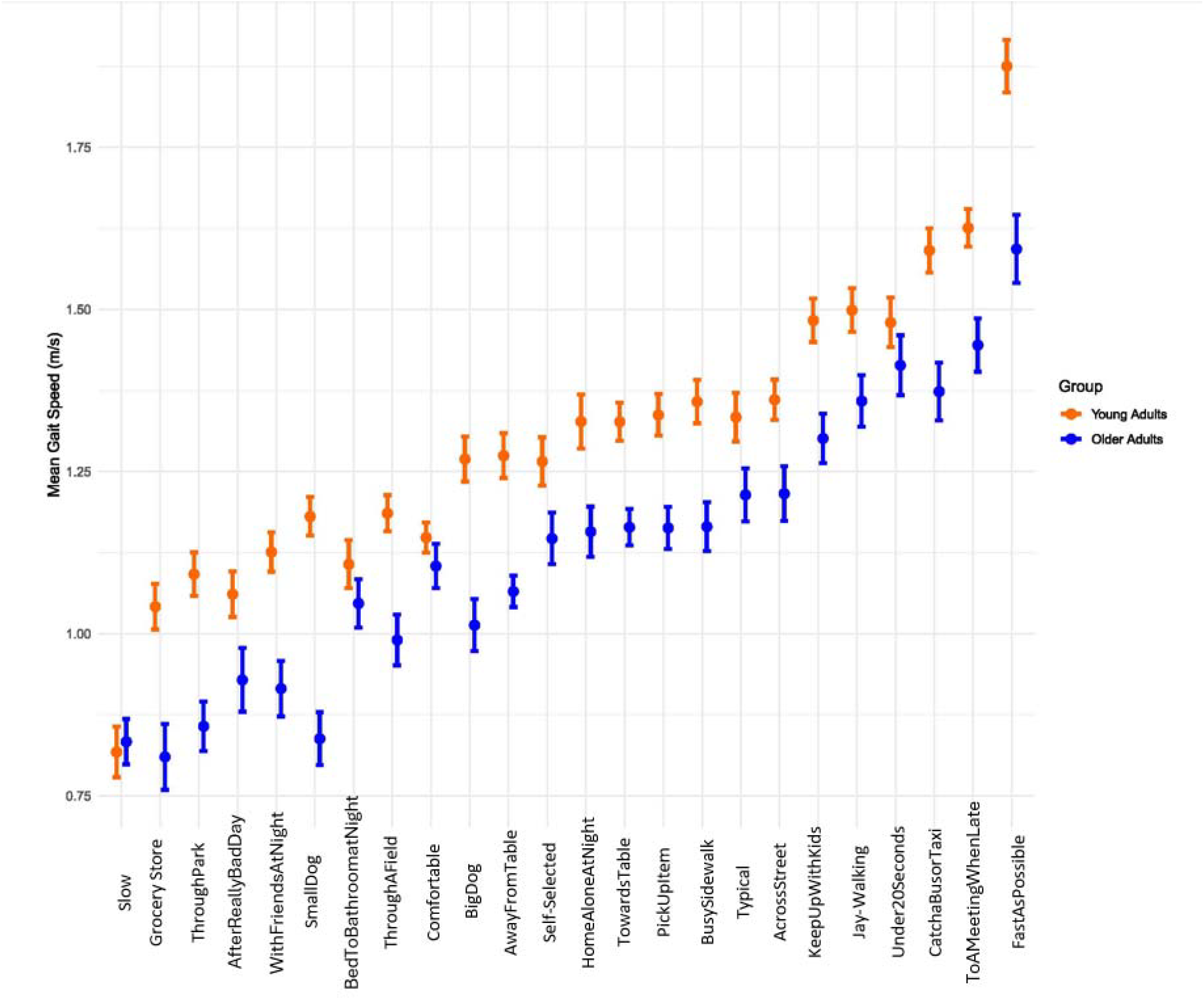
Distribution of speed by prompt ordered from slowest to fastest median speed, separated by group.

### Primary Aim 1: Variance Within Age Groups

Our first primary aim was to determine whether instructions explain similar amounts of variance within young and older adult groups when analyzed separately. We addressed this question through two complementary analytical approaches to ensure robustness of our findings.

When analyzing each age group as separate datasets, we calculated intraclass correlation coefficients (ICC) to determine the proportion of variance attributable to different walking prompts versus individual differences. This analysis revealed that instructions explained 55.7% of gait speed variance in young adults and 56.9% of gait speed variance in older adults (Table 2). To corroborate these findings, we conducted a second analysis using group-specific random effects modeling within a single mixed-effects framework. This approach yielded nearly identical results, with instructions explaining 55.2% of gait speed variance in young adults and 57.2% of gait speed variance in older adults.

**Table 2.** Variance components by analysis type.

| Analysis Method | Young Adults ICC | Older Adults ICC | Combined Model |
| --- | --- | --- | --- |
| Separate datasets | 55.7% | 56.9% | N/A |
| Group-specific random effects | 55.2% | 57.2% | N/A |
| Cross-classified model | N/A | N/A | 76.1% |
| Between-subject only | N/A | N/A | 25.3% |

Statistical comparison of these group-specific ICCs through bootstrapping methods (5000 simulations) revealed no significant difference between age groups (p = .85). This finding indicates that the degree to which instructions influence gait speed selection remains remarkably consistent across the lifespan, with verbal cues accounting for approximately 56% of the variance in walking behavior within both young and older adult populations. The consistency of this finding across two different analytical approaches strengthens confidence in this conclusion and suggests that instruction-based variability represents a fundamental characteristic of human gait control that is preserved with aging.

### Primary Aim 2: Age Differences in Response to Specific Instructions

Our second primary aim was to examine whether young and older adults respond differently to the same walking instructions, despite the similar proportions of variance explained within each group. This analysis required a systematic examination of variance components and interaction effects.

We first estimated an unconditional model with only a random intercept for subject to establish baseline variance decomposition. The ICC for this model was 0.253, indicating that 25.3% of gait speed variance is explained by between-subject differences alone (Table 3). Subsequently, we added a random intercept for instruction type in addition to the random intercept for subject, creating a cross-classified model that explained 76.1% of total variance when both random intercepts were included simultaneously. This substantial increase in explained variance (from 25.3% to 76.1%) demonstrates the powerful influence of instruction type on gait speed selection. Model comparison confirmed that the cross-classified model provided significantly better fit than the subject-only model (χ^2^(23) = 1280.9, p < 0.001), with superior Akaike information criterion (−932.51 vs 346.34) and Bayesian information criterion (−906.56 vs 367.10) values.

**Table 3.** Multilevel Models Comparison.

| Model Component | Model 1: Subject-Only | Model 2: Subject + Instruction |
| --- | --- | --- |
| Fixed Effects |  |  |
| Intercept | 1.21* (0.09) | 1.21* (0.10) |
| Random Effects Variance |  |  |
| Subject | 0.013 | 0.020 |
| Instruction | -- | 0.050 |
| Residual | 0.070 | 0.020 |
| Model Fit |  |  |
| AIC | 346.34 | -932.51 |
| BIC | 367.10 | -906.56 |
| Log-likelihood | -167.17 | 471.26 |
| Model Comparison |  |  |
| $\chi^2$ (df) | -- | 1280.9 (23)*** |
| Variance Explained |  |  |
| ICC (Subject) | 25.3% | 28.6% |
| ICC (Instruction) | -- | 47.6% |
| Total explained | 25.3% | 76.1% |

**\*p < 0.05**

To specifically test whether age groups respond differently to individual instructions, we fitted a model including an interaction between age group and instruction type. This analysis revealed a significant age-instruction interaction (χ^2^ = 76.84, df = 23, p < .001), indicating that while instructions explain similar proportions of variance within each age group, they simultaneously create systematic differences in how each age group responds to the same verbal prompts. The higher variance explained in the combined model (76.1%) compared to within-group analyses (~56%) reveals a fascinating paradox: the variance of instruction influence remains consistent across the lifespan, yet the magnitude of that influence changes systematically with age.

### Detailed Interaction Analysis

The interaction model produced β coefficients that quantified the magnitude of age-related differences for each specific instruction (Table 4). These coefficients represent how much slower (in m/s) older adults walked compared to young adults for each instruction, after accounting for individual variation through random subject effects.

**Table 4.** Age-Instruction Interaction Coefficients.

| Instruction | $\beta$ Coefficient | p-value |
| --- | --- | --- |
| Walk as if walking a small dog | -0.316 | 7.68e-09 |
| Walk as fast as possible | -0.241 | 1.19e-05 |
| Walk as if walking in grocery store | -0.209 | 0.000134 |
| Walk home with friends at night | -0.203 | 0.000196 |
| Walk as if walking a big dog | -0.203 | 0.000190 |
| Walk away from the table | -0.191 | 0.000438 |
| Walk as if walking through a park | -0.188 | 0.000550 |
| Walk as if walking through a field | -0.185 | 0.000744 |
| Walk to catch a bus or taxi | -0.176 | 0.001194 |
| Walk to pick item up off table | -0.165 | 0.002765 |
| Walk at speed on busy sidewalk | -0.159 | 0.003819 |
| Walk towards the table | -0.152 | 0.005116 |
| Walk trying to keep up with kids | -0.154 | 0.005167 |
| Walk home alone at night | -0.145 | 0.007605 |
| Walk to meeting when late | -0.145 | 0.008661 |
| Walk across the street | -0.135 | 0.013885 |
| Walk home after bad day | -0.125 | 0.021179 |
| Walk at your typical speed | -0.124 | 0.023674 |
| Walk as if jay-walking | -0.120 | 0.027591 |
| Walk at self-selected speed | -0.103 | 0.057979 |
| Walk in under 20 seconds | -0.060 | 0.269867 |
| Walk at comfortable speed | -0.037 | 0.491638 |
| Walk bed to bathroom at night | -0.027 | 0.625332 |

Analysis of the interaction coefficients revealed clear patterns in the types of instructions that elicited the largest age-related differences. Instructions involving multiple cognitive components or time constraints showed the most pronounced between-group differences. Specifically, ‘walking a small dog’ produced the largest age-related difference (β = −0.316, p < 0.001), followed by ‘walk as fast as possible’ (β = −0.241, p < 0.001). These findings suggest that older adults may be particularly conservative in responding to complex or ambiguous verbal instructions, possibly reflecting enhanced attention to safety or different interpretation of instruction ambiguity.

Conversely, simple, direct clinical instructions showed minimal and statistically non-significant differences between age groups. Most notably, ‘walk from bed to bathroom at night’ (β = −0.027, p = .625), ‘walk at a comfortable speed’ (β = −0.037, p = .492), and ‘walk in under 20 seconds’ (β = −0.060, p = .270) produced the smallest age-related differences. These patterns suggest that straightforward, unambiguous instructions produce more consistent responses across age groups.

The consistently negative interaction coefficients demonstrate that older adults showed a systematically smaller range of adjustments to their walking speed across different instructions compared to young adults. This pattern suggests older adults may employ a more cautious approach to modulating their gait speed, potentially reflecting age-related changes in balance confidence, motor control strategies, or risk assessment. The magnitude of these differences varied considerably across instructions, with complex instructions showing effect sizes of 0.24 to 0.32 m/s, while simple instructions showed minimal differences of 0.03 to 0.06 m/s.

### Variance Homogeneity Analysis

Despite systematic differences in mean gait speed responses between age groups, analysis of variance homogeneity revealed remarkably consistent variability patterns within each age group across most instructions. Levene’s tests conducted for each individual instruction showed similar variance between age groups across 23 of 24 instructions, with only one instruction (“Walk away from the table”) showing significantly different variance between groups (p = .039). This finding suggests that while absolute speeds may differ between age groups for specific instructions, the relative consistency of response within each group remains stable across the lifespan.

Figure 3 illustrates the variance patterns by instruction for both age groups, demonstrating that both young and older adults show similar patterns of variability across different walking prompts. This consistency in variance patterns has important implications for both research methodology and clinical practice, indicating that reliable measurements can be obtained within age groups even though the absolute speeds may differ between groups. The preservation of within-group consistency suggests that instruction-based gait assessment protocols maintain their measurement properties across different age populations.

**Fig. 3.**
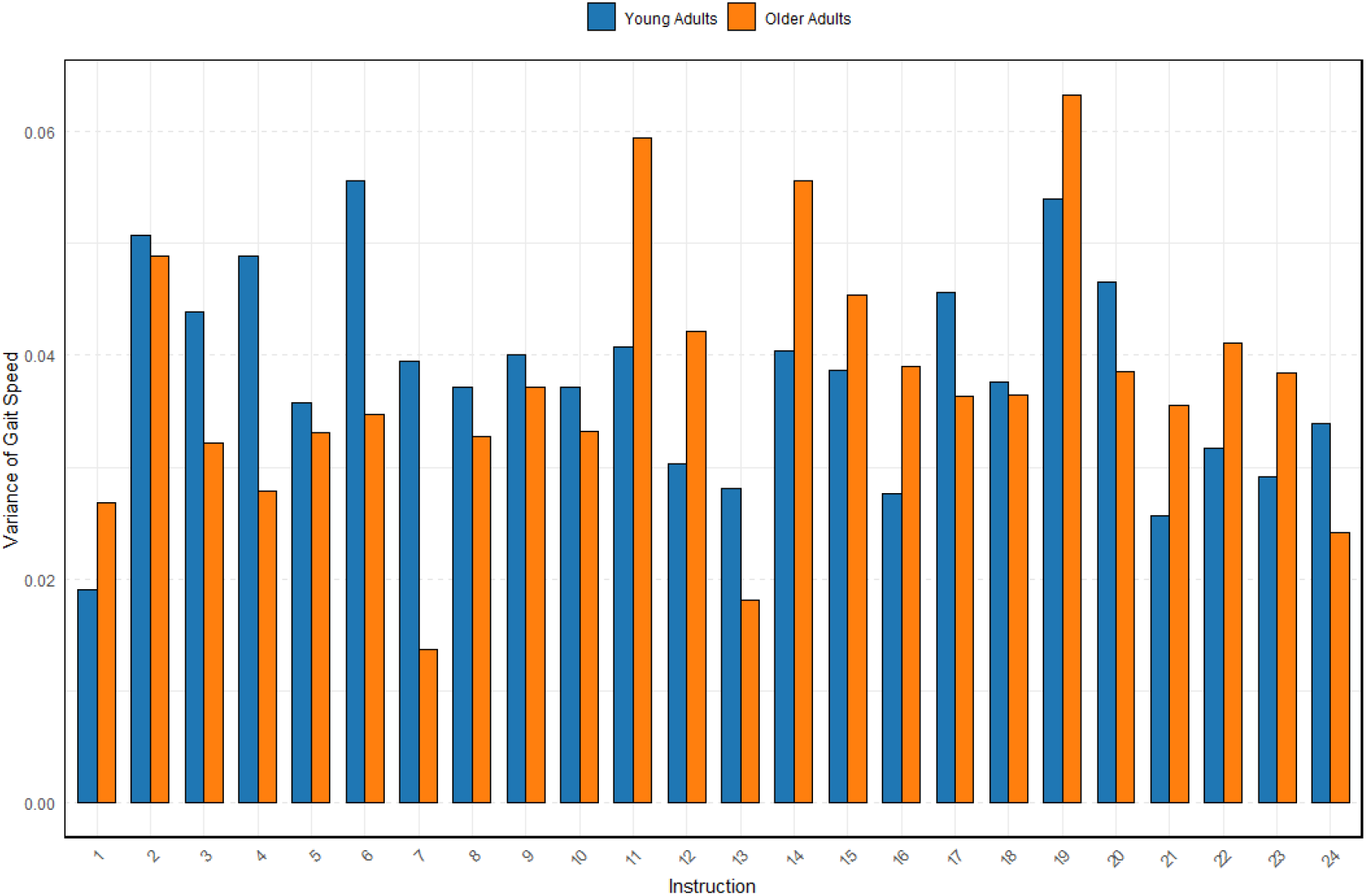
Variance in gait speed by instruction number for young adults (blue bars) and older adults (orange bars). The y-axis shows the variance of gait speed in m/s^2^, and the x-axis shows the 24 different instructions. Higher bars indicate greater variability in gait speed for that instruction within each age group.

### Summary of Statistical Findings

Three key statistical findings emerged from these analyses. First, instructions explained approximately 56% of gait speed variance within both young adults (55.7%) and older adults (56.9%), with no significant difference between age groups (p = .85). Second, when analyzing both age groups together, the cross-classified model including random intercepts for both subjects and instructions explained 76.1% of total variance, compared to 25.3% when including only subject effects. Third, a significant age-instruction interaction (χ^2^ = 76.84, df = 23, p < .001) indicated that age groups responded differently to specific instructions, with complex instructions producing larger between-group differences (β: −0.24 to −0.32 m/s) than simple instructions (β: −0.03 to −0.06 m/s).

## Discussion

The primary aims of this study were to investigate how verbal instructions influence gait speed in both young and older adults, and to determine whether age-related differences exist in response to various walking prompts. Our findings revealed two key outcomes:

1. When analyzed separately, instructions explained similar proportions of variance within older adults (56.9%) and young adults (55.7%) with no significant difference between groups.
2. Instruction type significantly explained 76.1% of gait speed variance when analyzing both age groups together, suggesting different age-specific response patterns to instructions.

The consistency in variance explained by instructions within each age group (approximately 56%) is remarkable, suggesting that the degree to which instructions influence gait selection remains important across the lifespan, even as absolute speeds differ. This finding indicates that verbal cues maintain a similarly powerful effect on walking behavior regardless of age, with instructions accounting for more than half of the variance in how people walk within their respective age cohorts. However, the higher variance explained in the combined model (76.1%) reveals a fascinating paradox: while instructions explain similar proportions of variance within each age group (~56%), they simultaneously create systematic differences in how each age group responds. This pattern suggests that the *variance* of instruction influence remains consistent across the lifespan, yet the *magnitude* of that influence changes with age.

This dual effect is confirmed by our interaction analysis, which revealed significant interactions between age group and instruction type (χ^2^(23) = 76.84, p < .001). Older and younger adults interpret and respond differently to the same verbal prompts, despite instructions explaining similar proportions of variance within each group. The consistently negative interaction coefficients demonstrate that older adults show a smaller range of adjustments to their walking speed across different instructions compared to young adults. This pattern suggests older adults may employ a more cautious approach to modulating their gait speed, potentially reflecting age-related changes in balance confidence or motor control strategies.

Instructions that elicited the largest age-related differences typically involved more complex scenarios or time-pressured situations, with interaction coefficients exceeding −0.25. This suggests that older adults may be particularly conservative in adjusting their gait speed when tasks involve multiple components or environmental challenges, possibly reflecting enhanced attention to safety or reduced confidence when given complex walking instructions. Conversely, instructions involving simple, straightforward walking tasks showed minimal age-related differences, suggesting these prompts may be more suitable for standardized assessment across age groups. Despite these age-related differences in response to instructions, we found remarkably consistent variance between age groups across most individual instructions, with only one instruction showing significantly different variance between groups. This suggests that while absolute speeds may differ between age groups, the relative consistency of response within each group remains stable. This finding has important implications for both research methodology and clinical practice, indicating that reliable measurements can be obtained within age groups even though the absolute speeds may differ between groups.

Our findings complement our previous research [1] by demonstrating that while instructions explain similar proportions of variance within each age group (~56%), there are clear age-dependent differences in how individuals respond to specific instructions. This nuance represents an important advancement in our understanding of gait assessment across the lifespan. The consistent pattern of more conservative gait speed adjustments in older adults, particularly for complex instructions, helps explain why 76.1% of variance is accounted for in the combined model despite explaining only ~56% within each age group separately. This additional variance is captured by certain instructions eliciting the systematic age-related differences in response patterns, while other instructions do not. These findings should be interpreted considering methodological differences from our previous 2022 study [1]. While the previous study used an instrumented walkway for speed measurement, the current study employed wearable APDM sensors. The similar proportion of variance explained by instructions in young adults (61% in 2022 vs ~56% in current study) suggests that instruction effects are robust across measurement methodologies, strengthening confidence in the generalizability of these findings across different gait assessment approaches.

The phenomenon of more conservative speed adjustments in older adults aligns with established theories of aging and motor control [4, 5, 6, 7]. These differences likely reflect an adaptive strategy that prioritizes stability and safety over speed, particularly during instructions involving complex cognitive demands or perceived risk. Instructions that showed minimal between-group differences in response patterns may represent ideal candidates for standardized clinical assessments, as they produce more comparable results across age groups. When the goal is to measure age-related changes in gait parameters, clinicians and researchers should ensure the instructions being used are demanding enough to elicit these differences.

## Limitations

Several limitations should be acknowledged when interpreting these findings. Our sample consisted of healthy adults from a university community, potentially limiting generalizability to more wide-ranging populations with varying educational backgrounds, cognitive status, or cultural contexts that might influence instruction interpretation. Additionally, while our protocol maintained consistent instruction delivery, real-world clinical assessments may involve variations in tone, emphasis, or contextual cues that could further influence gait selection. Future studies should examine whether the observed patterns of instruction-based variance persist in populations with cognitive impairment, where verbal processing deficits might alter the instruction-gait relationship. Extending this paradigm to clinical populations with specific gait pathologies could determine whether disease processes modify the relationship between verbal cues and motor execution. Longitudinal investigations tracking how instruction-based variability changes throughout the aging process within individuals would be particularly valuable, potentially revealing when and how these age-related differences in instruction response emerge. Finally, integrating cognitive assessments with these gait protocols could elucidate the executive function components most closely associated with instruction processing during gait planning.

## Conclusions

The identification of instructions producing minimal between-group differences has substantial clinical utility for standardizing gait assessment protocols. Instructions such as ‘Walk at a comfortable speed’ showed the smallest age-related differences, suggesting these prompts may provide more comparable assessments across age groups. Clinicians conducting gait assessments across various age populations might prioritize these straightforward instructions to minimize age-related interpretation differences and improve measurement consistency. Conversely, when the goal is to detect age-related changes in gait adaptation or instruction processing, more complex or ambiguous instructions may be more revealing of functional differences. This selective approach to instruction selection would enhance the specificity and sensitivity of gait assessments, allowing clinicians to either minimize or highlight age-related differences depending on their assessment objectives. Standardizing instruction protocols based on these findings could reduce measurement variability in multi-site clinical trials and improve the reliability of longitudinal assessments spanning significant time periods.

## References

[1] S. A. Brinkerhoff, W. M. Murrah, Z. Hutchison, M. Miller, and J. A. Roper, “Words matter: instructions dictate ‘self-selected’ walking speed in young adults,” Gait & Posture, vol. 95, pp. 223–226, Jun. 2022, doi: 10.1016/j.gaitpost.2019.07.379.

[2] S. Fritz and M. Lusardi, “White paper: ‘walking speed: the sixth vital sign,’” J Geriatr Phys Ther, vol. 32, no. 2, pp. 46–49, 2009.

[3] R. W. Bohannon and A. Williams Andrews, “Normal walking speed: a descriptive meta-analysis,” Physiotherapy, vol. 97, no. 3, pp. 182–189, Sep. 2011, doi: 10.1016/j.physio.2010.12.004.

[4] R. Morris, S. Stuart, G. McBarron, P. C. Fino, M. Mancini, and C. Curtze, “Validity of MobilityLab (version 2) for gait assessment in young adults, older adults and Parkinson’s disease,” Physiol Meas, vol. 40, no. 9, p. 095003, Sep. 2019, doi: 10.1088/1361-6579/ab4023.

[5] B. T. Cleland, T. Alex, and S. Madhavan, “Concurrent validity of walking speed measured by a wearable sensor and a stopwatch during the 10-meter walk test in individuals with stroke,” Gait Posture, vol. 107, pp. 61–66, Jan. 2024, doi: 10.1016/j.gaitpost.2023.09.012.

[6] M. Mancini and F. B. Horak, “Potential of APDM Mobility Lab for the monitoring of the progression of Parkinson’s disease,” Expert Rev Med Devices, vol. 13, no. 5, pp. 455–462, May 2016, doi: 10.1586/17434440.2016.1153421.

[7] X. Fang, C. Liu, and Z. Jiang, “Reference values of gait using APDM movement monitoring inertial sensor system,” Royal Society Open Science, vol. 5, no. 1, p. 170818, Jan. 2018, doi: 10.1098/rsos.170818.

[8] J. L. Allen and J. R. Franz, “The motor repertoire of older adult fallers may constrain their response to balance perturbations,” J Neurophysiol, vol. 120, no. 5, pp. 2368–2378, Nov. 2018, doi: 10.1152/jn.00302.2018.

[9] K. A. Boyer et al., “Age-related changes in gait biomechanics and their impact on the metabolic cost of walking: Report from a National Institute on Aging workshop,” Experimental Gerontology, vol. 173, p. 112102, Mar. 2023, doi: 10.1016/j.exger.2023.112102.

[10] S. A. Brinkerhoff, W. M. Murrah, and J. A. Roper, “The relationship between gait speed and mediolateral stability depends on a person’s preferred speed,” Sci Rep, vol. 13, no. 1, p. 6056, Apr. 2023, doi: 10.1038/s41598-023-32948-z.

[11] J. L. Campos, U. Marusic, and J. R. Mahoney, “Editorial: The intersection of cognitive, motor, and sensory processing in aging: Links to functional outcomes, Volume I,” Frontiers in Aging Neuroscience, vol. 14, 2022, Accessed: Sep. 20, 2023. [Online]. Available: https://www.frontiersin.org/articles/10.3389/fnagi.2022.1009532

